# Canonical Wnt induction by OTULIN prevents keratinocyte death and skin inflammation

**DOI:** 10.1101/2025.01.08.631848

**Authors:** Kim Lecomte, Pieter Hertens, Annagiada Toniolo, Fleur Boone, Maarten Ciers, Lien Verboom, Katrien Staes, Geert van Loo, Esther Hoste

## Abstract

Loss-of-function mutations in the human *OTULIN* gene, encoding a deubiquitinase with exclusive specificity for linear ubiquitin chains, cause a severe multi-organ autoinflammatory condition involving the skin^1,2^. Mice lacking OTULIN selectively in keratinocytes develop inflamed skin lesions that progress into squamous tumours, a phenotype driven by excessive TNF-induced cell death^3,4^. Previous studies suggested a role for OTULIN in mediating Wnt signalling during development^5^, but the physiological relevance of this association is unknown. Here, we show that OTULIN promotes Wnt signalling in keratinocytes by regulating the linear ubiquitination status of β-catenin. Stabilisation of β-catenin in OTULIN-deficient keratinocytes prevents progressive skin inflammation in prophylactic and therapeutic settings by blocking keratinocyte death. We demonstrate that linearly ubiquitinated β-catenin accumulates in OTULIN-deficient keratinocytes, promoting its ubiquitination with K48 chains and subsequent proteasomal degradation. Reduced Wnt signalling in OTULIN-deficient keratinocytes leads to degradation of TCF3, an essential survival factor for keratinocytes^6^. Collectively, our data identify OTULIN’s linear deubiquitination activity as a key regulator of epithelial cell viability, not only by preventing cell death downstream of TNF, but also by promoting canonical Wnt signalling.

---

Keratinocytes (KCs), the epithelial cells of the skin, constitute a crucial structural and immunological barrier protecting the organism from dehydration and extrinsic hazards^7^. The survival of KCs under external insults critically depends on their precise regulation of inflammatory signalling cascades, which are predominantly mediated through specific and fast post-translational modifications of signalling proteins, ultimately triggering downstream transcriptional responses^8^. Ubiquitination is such a post-translational modification important for orchestrating cellular responses in inflammation. Linear ubiquitin (M1) chains are head-to-tail linked ubiquitin moieties important for assembling protein complexes downstream of tumour necrosis factor (TNF), interleukin-1β (IL-1β) and Toll-like receptors^9–12^. M1 chains are added to substrates by the linear ubiquitin chain assembly complex (LUBAC), consisting of haem-oxidised IRP2 ubiquitin ligase-1 (HOIL-1), HOIL interacting protein-1 (HOIP) and Shank Associated RH Domain Interacting protein (SHARPIN)^13^. These M1 Ubq chains can be removed from substrates by the deubiquitinases (DUBs) OTU DUB with linear linkage specificity (OTULIN, encoded by the gene *Fam105b*, also known as *gumby*) and Cylindromatosis (CYLD). OTULIN is the only DUB with exclusive specificity for M1 ubiquitin chains, while CYLD is also capable of removing K63 ubiquitin chains^14–16^. Importantly, OTULIN prevents LUBAC autoubiquitination and degradation, stabilising LUBAC in specific cell-types, including keratinocytes, thereby mediating NF-κB signalling and preventing cell death^1,3,4,17,18^.

Patients harbouring loss-of-function mutations in *OTULIN* develop a severe, often lethal, disease with skin involvement, termed ORAS (OTULIN-related auto-inflammatory syndrome), and are successfully treated with TNF-blocking therapies^1,2^. Genetic deletion of *Otulin* or expression of a catalytic inactive OTULIN mutant in mice induces embryonic lethality due to excessive TNFR1-mediated cell death^1,5,17^. We and others previously showed that mice lacking OTULIN selectively in KCs develop severe skin inflammation driven by TNFR1 signalling^3,4^. *Rivkin* et al. identified the *Otulin/gumby* gene as causative for the recessive, embryonically lethal gumby mouse mutant and suggested a critical role for the gumby-LUBAC axis in regulating canonical Wnt signalling^5^. However, *in vivo* evidence of such a role is currently lacking. Wnt signalling pathways regulate a diverse range of cellular processes, including cell fate determination, proliferation and migration^19^. Wnt is crucial for timely adaption of embryonic and adult stem cell behaviour and induces hair follicle stem cell proliferation in skin, but also controls commitment and differentiation of other KCs^20,21^. Canonical Wnt signalling depends on β-catenin and in absence of Wnt ligands, β−catenin is bound by a destruction complex comprising glycogen synthase-kinase-3β (GSK3β), axis inhibition 2 (AXIN2), adenomatous polyposis coli (APC) and casein kinase 1α (CK1α), resulting in β-catenin phosphorylation. Phosphorylated β−catenin is subsequently recognized by β-transducin repeat containing E3 ubiquitin protein ligase (β-TrCP), which catalyses the addition of K48-linked ubiquitin chains, thereby targeting β−catenin for proteasomal degradation^22^. Upon binding of Wnt ligands to its receptors, Frizzled (Fzd) and its co-receptor low-density lipoprotein receptor-related protein 5/6 (LRP5/6), the adaptor Dishevelled2 (DVL2) is recruited to the membrane and binds AXIN2, thereby disrupting the destruction complex. This disruption allows β−catenin to accumulate in the cytosol and translocate to the nucleus, where it activates transcription of Wnt target genes by forming a complex with members of the T cell factor (TCF)/lymphoid enhancer-binding factor (LEF) family of transcription factors^23^. The E3 ubiquitin ligases RNF43 and ZNRF3 function as Wnt repressors by catalysing K48-linked ubiquitination, which triggers the turnover or endocytosis of Fzd receptors^24,25^, while β-catenin degradation is predominantly mediated by the SCF E3 ligase complex containing β-TRCP^22,26^. Interestingly, overexpression studies in cancer cell lines suggested that linear ubiquitination of β-catenin promotes its K48-linked ubiquitination and degradation in response to DNA damage, a process that is counteracted by OTULIN^27^.

In this study, we elucidate the critical *in vivo* importance of OTULIN in regulating epithelial canonical Wnt signalling. We demonstrate that stabilisation of β-catenin and expression of the Wnt target genes TCF3/4 depend on OTULIN’s DUB activity, thereby protecting KCs from TNFR1-induced cell death and severe skin inflammation in mice. Notably, stabilising β-catenin in KC-selective OTULIN-deficient mice prevents or ameliorates dermatitis in both prophylactic and therapeutic settings. These findings unveil important new insights into how morphogen signalling intersects with cell death pathways to safeguard epithelial cell survival.

## Results

### Wnt signalling is downregulated in OTULIN-deficient keratinocytes

OTULIN was initially characterised as a modulator of canonical Wnt signalling^5^. When crossing OTULIN-deficient mice with *TOPGAL* reporter mice, a role for OTULIN in promoting Wnt signalling during embryonic angiogenesis was suggested^5^. To analyse whether OTULIN and linear (de)ubiquitination modulate Wnt signalling in adult tissues, we investigated Wnt signalling in skin of mice with OTULIN-deficient KCs (Δ^Ker^OTULIN mice). We previously demonstrated that Δ^Ker^OTULIN mice develop inflamed skin lesions that progress into well-differentiated squamous carcinomas^3,4^. Analysis of our published single-cell RNA-sequencing (scRNAseq) dataset comparing total skin of wild-type (Cre-negative OTULIN^fl/fl^) and Δ^Ker^OTULIN mice, showed downregulation of several Wnt mediators and target genes in OTULIN-deficient KCs relative to control KCs (Figure 1a, Supplementary Figure 1a). Quantitative PCR analysis of epidermal tail lysates confirmed the downregulation of Wnt target genes in Δ^Ker^OTULIN epidermis (Figure 1b). Expression levels of several members of the TCF/LEF family, key downstream mediators of canonical Wnt signalling, were also significantly reduced in Δ^Ker^OTULIN epidermis (Figure 1c). Wnt activation in adult KCs is most prominent during the growth phase of hair follicles (HFs), also referred to as anagen, culminating in nuclear expression of LEF1^21,28^. When probing HF cell populations (outer and inner layer, bulge and upper HF) of our scRNAseq data for Wnt target gene expression, the downregulation of Wnt signalling in OTULIN-deficient KCs was even more pronounced than in the total KC population (Figure 1d). Immunoblotting for non-phosphorylated β-catenin, indicative of stabilised β-catenin and active Wnt signalling, confirmed reduced Wnt activation in epidermal tail lysates from Δ^Ker^OTULIN relative to control mice (Figure 1e). Immunofluorescent staining of anagen skin indicated nuclear LEF1 in wild-type HFs, whereas anagen-like HFs in lesional Δ^Ker^OTULIN skin lacked nuclear LEF1 (Figure 1f). β-catenin also performs Wnt-independent roles at the membrane, therefore we assessed the levels of cytosolic β-catenin in OTULIN-deficient and control epidermis and observed higher levels in control epidermis, indicating that OTULIN promotes canonical Wnt signalling in KCs (Supplementary Figure 1b). Gene ontology (GO) enrichment analysis of genes markedly downregulated in OTULIN-deficient KCs relative to control KCs in our scRNAseq dataset confirmed that Wnt signalling is significantly downregulated in OTULIN-deficient KCs (Figure 1g). Together, these data demonstrate that OTULIN promotes canonical Wnt signalling in adult KCs.

**Figure 1.**
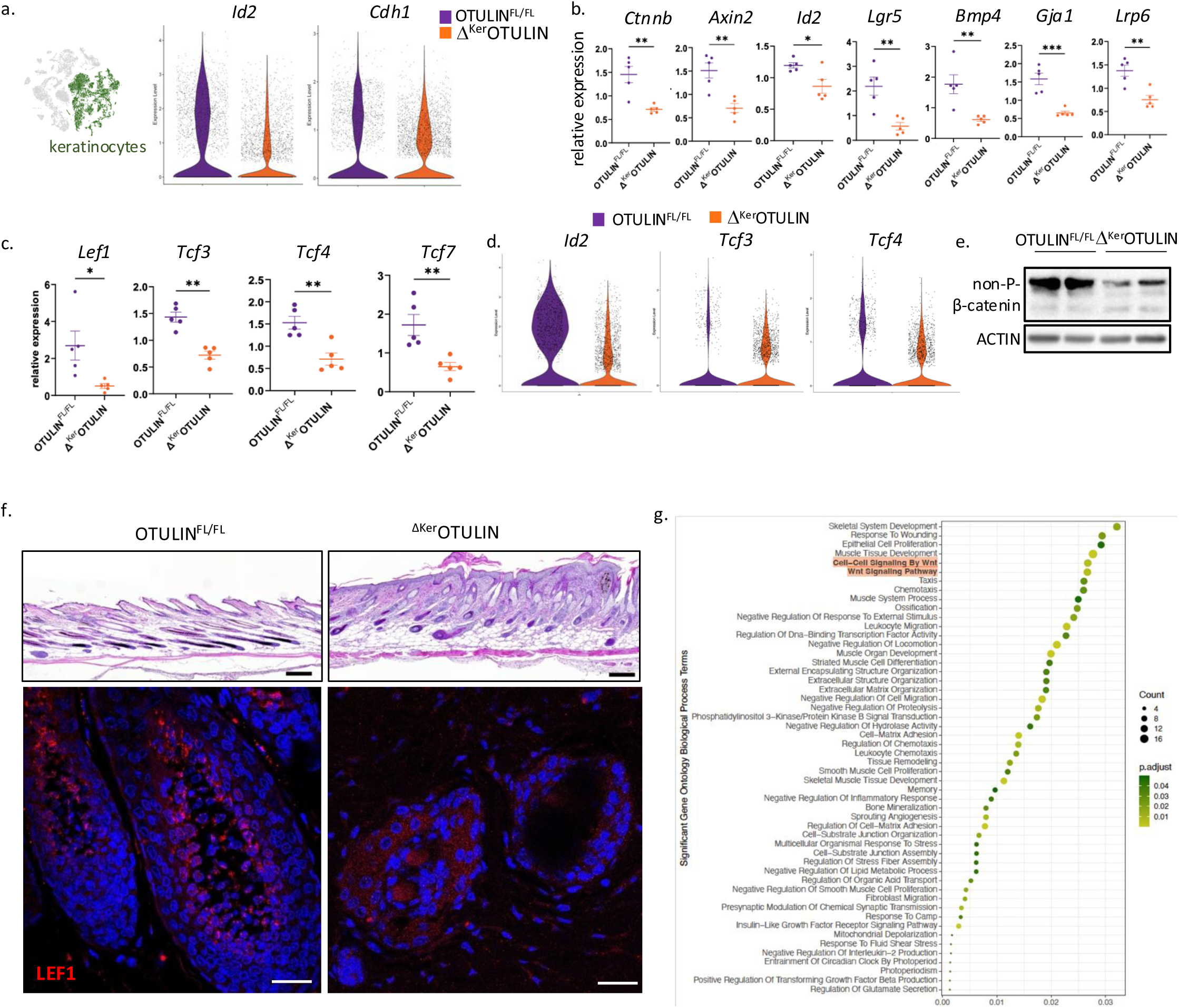
Wnt signalling is downregulated in OTULIN-deficient KCs. **a.** (left) UMAP clustering of scRNAseq data obtained from cells isolated from Δ^Ker^OTULIN and control skin^3^, highlighting the keratinocyte (KC) cluster. (right) Violin plots showing distribution of expression of the indicated Wnt target genes in scRNAseq data of KCs from OTULIN^FL/FL^ and Δ^Ker^OTULIN skin. **b.** Quantitative PCR (QPCR) analysis for the indicated Wnt target genes on epidermal tail lysates of OTULIN^FL/FL^ and Δ^Ker^OTULIN mice (age 8-11 weeks). Data are means ±SEM, ^∗^p< 0.05; ^∗∗^p< 0.01; ^∗∗∗^p< 0.001. **c.** QPCR analysis for the indicated Wnt effector genes on epidermal tail lysates of OTULIN^FL/FL^ and Δ^Ker^OTULIN mice. Data are means ± SEM, ^∗^p< 0.05; ^∗∗^p< 0.01; ^∗∗∗^p< 0.001. **d.** Violin plots showing distribution of expression of the indicated genes in scRNAseq data of hair follicle KCs from OTULIN^FL/FL^ and Δ^Ker^OTULIN skin. **e.** Immunoblotting for non-phosphorylated (active) β-catenin in epidermal tail lysates from OTULIN^FL/FL^ and Δ^Ker^OTULIN mice. α-actin is shown as loading control. **f.** Immunofluorescence images of LEF1 staining in OTULIN^FL/FL^ anagen skin and Δ^Ker^OTULIN lesional skin. Upper panels are H&E-stained skin sections of the same mice. **g.** Gene ontology (GO) enrichment analysis of genes significantly downregulated in scRNAseq data from OTULIN-deficient KCs relative to control KCs.

### Stabilisation of β-catenin prevents dermatitis and tumour formation in Δ^Ker^OTULIN mice

Canonical Wnt signalling is driven by cytosolic β-catenin accumulation and subsequent nuclear translocation, facilitating the interaction of β-catenin with TCF/LEF transcription factors to drive Wnt target gene expression. To assess whether inducing canonical Wnt signalling could compensate for the loss of OTULIN-driven Wnt promotion, we stabilised β-catenin in KCs of Δ^Ker^OTULIN mice. Given the crucial role of Wnt signalling in embryogenesis, we crossed Δ^Ker^OTULIN mice to K14ΔNβ-catenin-ER mice (hereafter referred to as ΔNβcat mice), which express a transgene lacking the N-terminal degradome of β-catenin under the control of a KC-selective promoter following postnatal tamoxifen induction^29^. Transgene induction in newborn Δ^Ker^OTULIN ΔNβcat pups at P0.5 and P1.5 provides complete protection from dermatitis in the tail skin. Approximately 20% of these mice also exhibit full protection from dermatitis in the back skin, even at old age (Supplementary Figure 2a). Other mice develop skin lesions, albeit significantly later than Δ^Ker^OTULIN mice (Figure 2a-c). Notably, all Δ^Ker^OTULIN ΔNβcat are protected from exophytic tumour formation (Supplementary Figure 2b). Time course measurements of transepidermal water loss (TEWL) indicated that Δ^Ker^OTULIN mice exhibit more severe epidermal barrier perturbation and increased epidermal thickness compared to Δ^Ker^OTULIN ΔNβcat littermates (Figure 2c, Supplementary Figure 2c). Clinical scoring of the mice further confirmed the significant improvement in skin pathology observed in Δ^Ker^OTULIN ΔNβcat relative to Δ^Ker^OTULIN mice (Figure 2d). Additionally, protection from skin inflammation in Δ^Ker^OTULIN ΔNβcat mice was evidenced by reduced levels of circulating interleukin-6 (IL-6) (Figure 2e).

**Figure 2.**
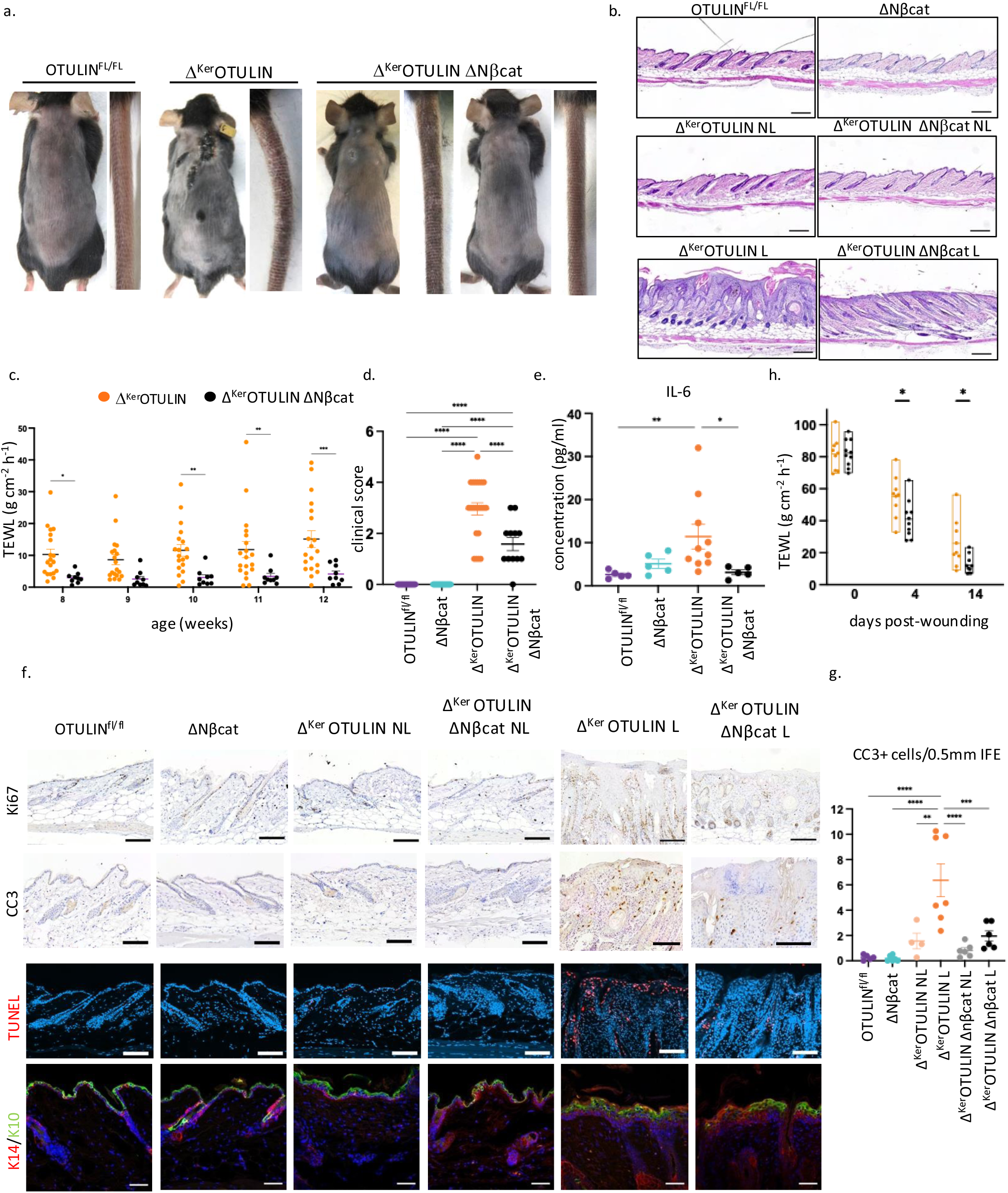
Stabilising β-catenin in KCs protects Δ^Ker^OTULIN skin from dermatitis and tumour formation. **a.** Representative images of the back skin and tail of 8 to 10-week old OTULIN^fl/fl^, Δ^Ker^OTULIN and Δ^Ker^OTULIN ΔNβ-cat mice. **b.** Representative images of H&E-stained skin sections of 10-week old mice of the indicated genotypes. Scale bar: 200 μm. NL: non-lesional; L: lesional. **c.** Time course TEWL measurements on lesional skin of Δ^Ker^OTULIN and Δ^Ker^OTULIN ΔNβcat mice (n> 4 mice per condition, multiple lesions per mouse were scored). Data are means ± SEM, ^∗^p< 0.05; ^∗∗^p< 0.01; ^∗∗∗^p< 0.001.****p< 0.0001). **d.** Clinical scoring (see Methods) of back and tail skin inflammation in mice of the indicated genotypes at week 14 of age. Data represent means ± SEM. (****p< 0.0001). **e.** Levels of IL-6 in serum of 10- to 14-week old mice of the indicated genotypes (n> 4 mice per genotype, *p< 0.05; ^∗∗^p< 0.01 Kruskal Wallis test). **f.** Immunostaining of skin sections from 10 to 14 week old mice of the indicated genotypes using antibodies against cleaved caspase 3 (CC3), Ki67, TUNEL, K10 and K14. Each staining was performed on 5 mice per condition. Scale bar: 200 μm. **g.** Quantification of the number of interfollicular epidermis (IFE) cells that stain positive for cleaved caspase-3 in skin sections from 10 to 14 week old mice of the indicated genotypes (OTULIN^FL/FL^ n=5; OTULIN^FL/FL^ ΔNβcat n=6; Δ^Ker^OTULIN NL n=4; Δ^Ker^OTULIN L n=6; Δ^Ker^OTULIN ΔNβ-cat NL n=6; Δ^Ker^OTULIN ΔNβcat L n=6). Data represent means ± SEM, *p< 0.05; **p< 0.01; ***p< 0.001. **h.** Wound healing kinetics by TEWL assessment after full-thickness wounding in Δ^Ker^OTULIN (n= 9) and Δ^Ker^OTULIN ΔNβcat (n= 10) mice (*p< 0.05; Two-way ANOVA with multiple comparisons).

Histomorphological analyses of the back skin revealed reduced epidermal hyperplasia in lesional skin of Δ^Ker^OTULIN ΔNβcat relative to Δ^Ker^OTULIN mice (Figure 2f, Supplementary Figure 2c), which was also confirmed by reduced Ki67 staining (Figure 2f). Given that skin pathology associated with defective linear (de)ubiquitination in KCs is driven by excessive cell death, we next questioned whether stabilisation of β-catenin protects OTULIN-deficient KCs from death. In lesional skin sections, a significant reduction in the number of apoptotic interfollicular epidermal cells is observed in Δ^Ker^OTULIN ΔNβcat skin compared to Δ^Ker^OTULIN skin as assessed by cleaved caspase-3 staining (Figure 2f, g). Similar reductions in cell death levels were observed using TUNEL (terminal deoxynucleotidyl transferase dUTP nick end labelling) staining, marking both apoptotic and necroptotic cells (Figure 2f). Immunostaining for keratin-10 (K10) and K14 revealed aberrant K14 expression in lesional skin of both Δ^Ker^OTULIN ΔNβcat and Δ^Ker^OTULIN mice (Figure 2f), indicating that KC differentiation is affected in lesional skin irrespective of the levels of β-catenin. OTULIN stabilises LUBAC components in various cell-types, including KCs^1,3,17^. One potential explanation for the significant improvement in the inflammatory phenotype of Δ^Ker^OTULIN mice upon β-catenin stabilisation is that LUBAC proteins might also be stabilised in these conditions. However, immunoblotting for SHARPIN and HOIP in epidermal lysates from Δ^Ker^OTULIN ΔNβcat and Δ^Ker^OTULIN mice showed a similar reduction of protein expression in both genotypes (Supplementary Figure 2d). Multiple reports suggest a role for Wnt signalling in skin wound healing^30,31^. When we subjected Δ^Ker^OTULIN and Δ^Ker^OTULIN ΔNβcat mice to full-thickness excisional skin wounding, we observed that Δ^Ker^OTULIN ΔNβcat mice close their wounds faster compared to Δ^Ker^OTULIN mice, as evidenced by accelerated epidermal barrier recovery (Figure 2h). Collectively, our findings demonstrate that skin inflammation driven by aberrant linear ubiquitination can be effectively mitigated *in vivo* by inducing canonical Wnt signalling.

### Therapeutic stabilisation of β-catenin ameliorates dermatitis in Δ^Ker^OTULIN mice

To assess whether inducing Wnt signalling can halt the progression of established skin lesions, Δ^Ker^OTULIN ΔNβcat and Δ^Ker^OTULIN mice were injected with tamoxifen at 3 and 4 weeks of age, a timepoint when lesions were already present. Macroscopic inspection of the skin revealed that inflammatory parameters, such as oedema and erythema, either remained static or improved over time in Δ^Ker^OTULIN ΔNβcat mice, in contrast to the progressive inflammation observed in Δ^Ker^OTULIN skin (Figure 3a, b). Clinical scoring confirmed the beneficial effect of Wnt induction in lesional Δ^Ker^OTULIN skin, reducing lesion progression and involvement of tail skin (Figure 3c). Time course analysis of TEWL in lesional skin of these mice further demonstrated that stabilisation of β-catenin prevents worsening barrier perturbation in Δ^Ker^OTULIN mice (Figure 3d). Moreover, cleaved caspase-3 and TUNEL staining indicated reduced cell death levels in lesional skin of Δ^Ker^OTULIN ΔNβ-cat mice relative to Δ^Ker^OTULIN mice (Figure 3e, f). These data show that promoting canonical Wnt signalling in KCs effectively halts the progression of skin inflammation caused by aberrant linear ubiquitination.

**Figure 3.**
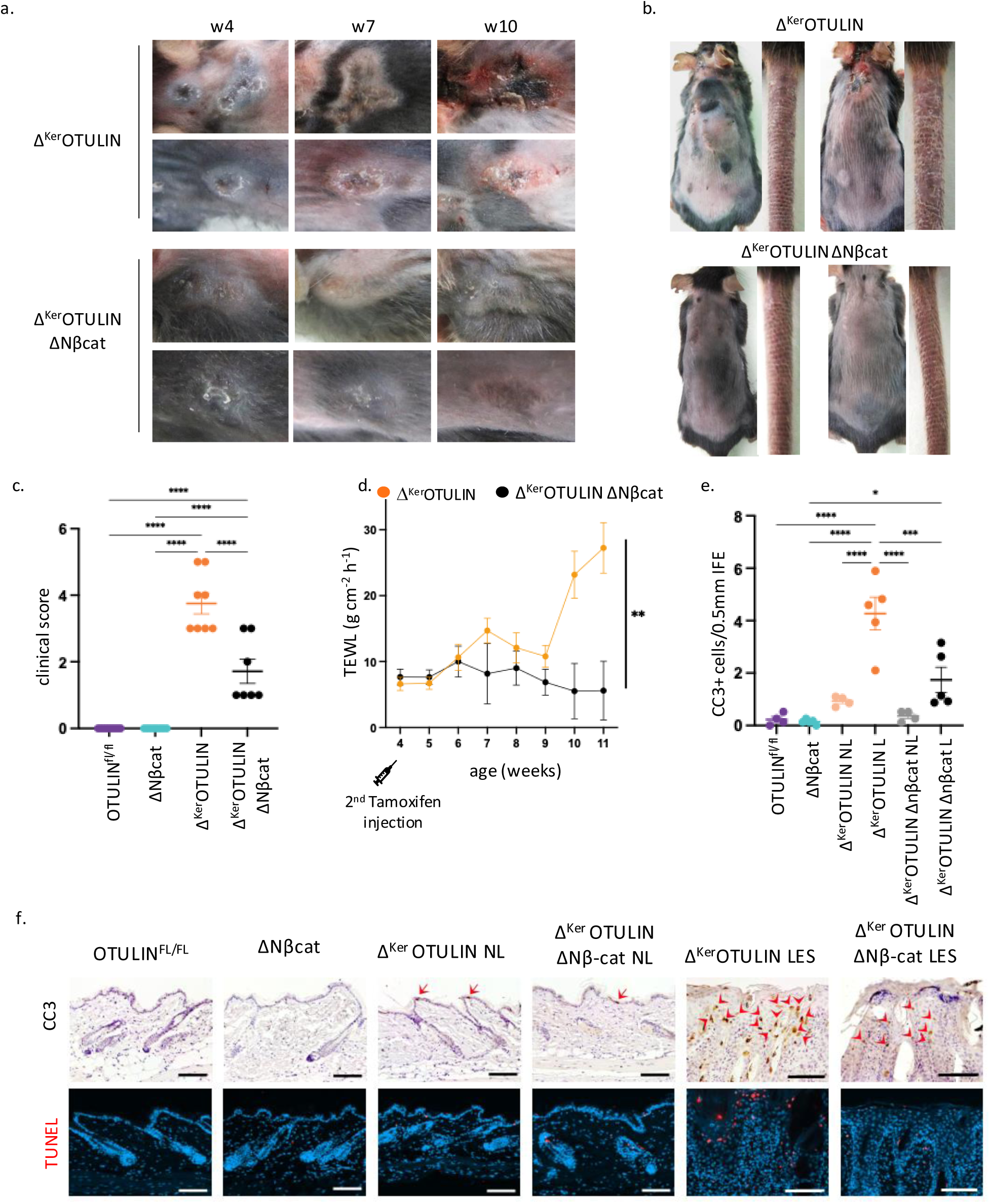
Stabilising β-catenin in established Δ^Ker^OTULIN skin lesions prevents progressive inflammation. **a.** Representative images of lesional back skin of Δ^Ker^OTULIN and Δ^Ker^OTULIN ΔNβ-cat mice at 4, 7 and 10 weeks of age. Mice were treated with tamoxifen at P21 and P28. **b.** Representative images of the back skin and tail of 8 to 10 week old Δ^Ker^OTULIN and Δ^Ker^OTULIN ΔNβ-cat mice treated with tamoxifen at P21 and P28. **c.** Clinical scoring of back and tail skin inflammation in mice of the indicated genotypes at week 14 of age. Data represent means ± SEM. (****p < 0.0001). **d.** Time course TEWL measurements on lesional skin of Δ^Ker^OTULIN and Δ^Ker^OTULIN ΔNβcat mice (n> 4 mice per condition, multiple lesions per mouse were scored, **p < 0.01). **e.** Quantification of the number of interfollicular epidermis (IFE) cells that stain for cleaved caspase-3 in skin sections from 10 to 14-week old mice of the indicated genotypes (OTULIN^FL/FL^ n=4; OTULIN^FL/FL^ ΔNβcat n=4; Δ^Ker^OTULIN NL n=4; Δ^Ker^OTULIN L n=5; Δ^Ker^OTULIN ΔNβ-cat NL n=4; Δ^Ker^OTULIN ΔNβcat L n=5). Data represent means ± SEM, *p< 0.5; ***p< 0.001; ****p< 0.0001. **f.** Immunostaining of skin sections from 10 to 14 week old mice of the indicated genotypes using antibody against cleaved caspase 3 (CC3) and TUNEL staining. Each staining was performed on n= 5 mice per condition. Scale bar: 200 μm. Arrows indicate apoptotic cells.

### The linear ubiquitin status of β-catenin controls its stability

Wnt/β-catenin activation has been previously shown to be regulated by ubiquitination^24–27^. To assess the linear ubiquitin status of β-catenin, we performed a specific pulldown using recombinant GST-UBAN (Ubq-binding domain in ABIN proteins and NEMO) on primary cultured KCs obtained from Δ^Ker^OTULIN (KO) and control OTULIN^fl/fl^ (WT) littermates. As previously reported, increased smearing of M1 chains was observed in OTULIN-deficient KCs^3^, both in pre-immunoprecipitation (PIP) and immunoprecipitated (IP) lysates (Figure 4a). Pulldown of UBAN containing proteins revealed linear ubiquitination of both total and unphosphorylated (active) β-catenin in primary mouse KCs (PMKs), which was slightly increased in absence of OTULIN (Figure 4a). Notably, the marked downregulation of active β-catenin observed in epidermal tail lysates from Δ^Ker^OTULIN mice (Figure 1e) was not sustained in these PMK cultures. To investigate the role of OTULIN in Wnt signalling in human KCs, we generated HaCaTs with strongly reduced OTULIN levels using OTULIN-targeting nanoblades (Supplementary Figure 3a, b). OTULIN knock-down (KD) HaCaT cells displayed increased sensitivity to TNF-induced cell death, consistent with our previous observations in OTULIN-deficient PMKs (Supplementary Figure 3c). Accumulation of M1 ubiquitin chains in OTULIN KD HaCaTs was evident in both PIP and IP lysates using UBAN beads following treatment of the cells with the proteasome inhibitor MG132 (Figure 4b). Pulldown of β-catenin with UBAN beads confirmed that β-catenin is linearly ubiquitinated in human KCs (Figure 4b). Further validation of the linear ubiquitin status of β-catenin in human KCs was performed through co-IP using a β-catenin-targeting antibody. Pulldown analysis revealed increased levels of linearly ubiquitinated β-catenin in OTULIN KD relative to control KCs, both in untreated and MG132-treated conditions. These experiments also indicate an interaction between β-catenin, OTULIN and the LUBAC protein SHARPIN (Figure 4c, Supplementary Figure 3d). Incubation of β-catenin IP lysates from OTULIN KD and control HaCaT cells with recombinant human OTULIN resulted in the collapse of the M1-linked ubiquitin smearing on both unphosphorylated and total pools of β-catenin, corroborating the linear ubiquitin status of β-catenin in untreated human KCs (Figure 4d). Co-IP studies using an anti-β-catenin antibody in PMKs isolated from Δ^Ker^OTULIN and control littermates confirmed the interaction between β-catenin and OTULIN in homeostatic conditions (Figure 4e). Additionally, pulldown of K48-linked ubiquitin proteins in PMK lysates from Δ^Ker^OTULIN and control skin indicated that, despite reduced total cytoplasmic β-catenin levels in PIP lysates from OTULIN-deficient KCs, the levels of K48-linked β-catenin were comparable in KCs of both genotypes in conditions of proteasomal inhibition (Figure 4f). Collectively, these findings demonstrate that OTULIN interacts with β-catenin in homeostatic KCs, modulating its linear ubiquitin status, which in turn influences its K48-linked ubiquitination and subsequent proteasomal degradation.

**Figure 4.**
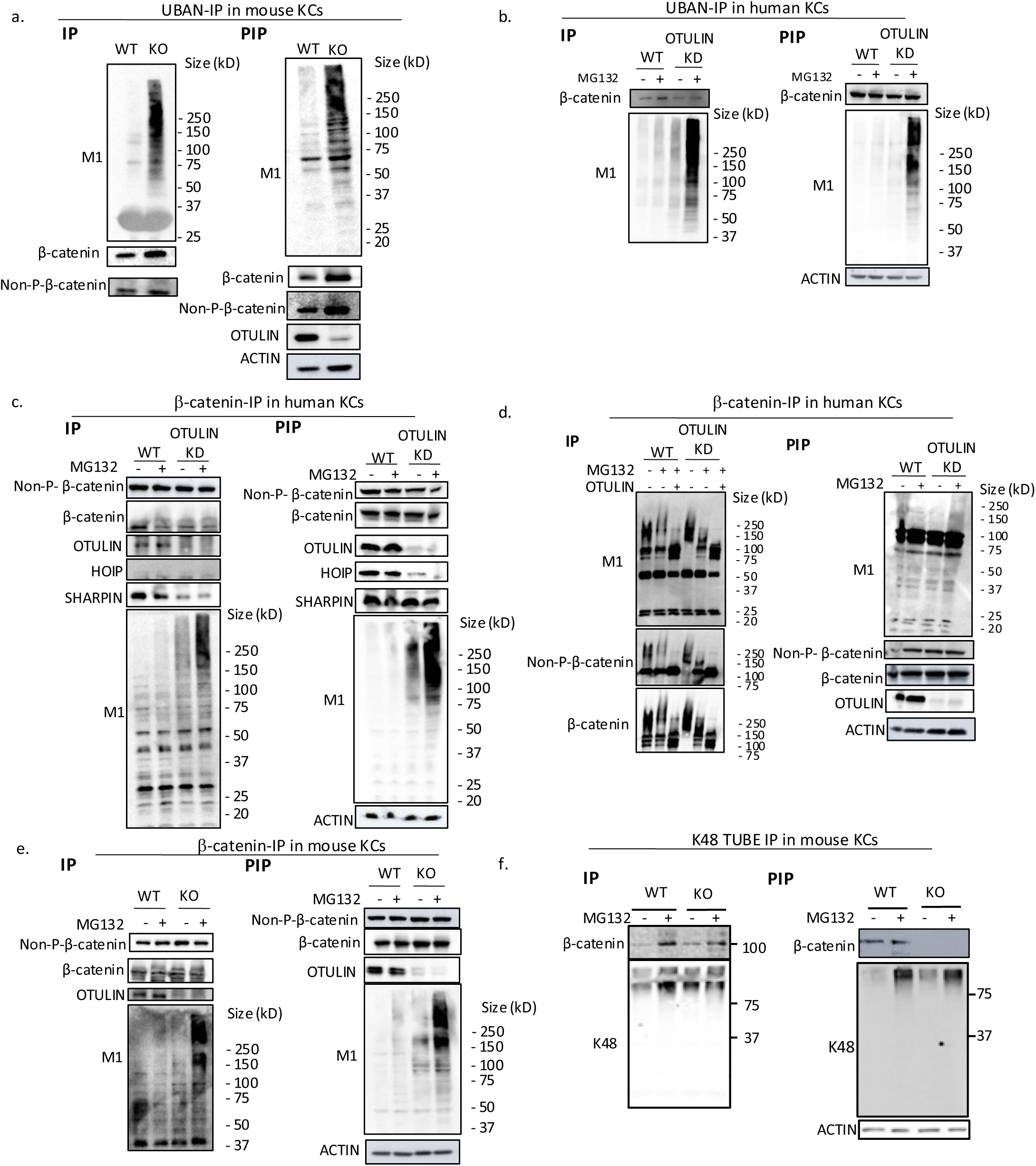
β-catenin is linearly ubiquitinated and coprecipitates with OTULIN in adult KCs. **a.** Linear ubiquitin pulldown by UBAN-immunoprecipitation (IP) on PMK cultures isolated from OTULIN^FL/FL^ and Δ^Ker^OTULIN mice followed by immunoblotting for M1 ubiquitin chains, non-phosphorylated (active) β-catenin and total β-catenin. Pre-immunoprecipitation (PIP) lysates were blotted for the same proteins and α-actin is shown as loading control. **b.** Linear ubiquitin pulldown on lysates from wild-type (WT) and OTULIN knock-down (KD) HaCaTs left untreated or treated with proteasome inhibitor MG132 for 16 hrs. α-actin is shown as loading control. **c.** Co-immunoprecipitation (IP) using an antibody against β-catenin on lysates from WT and OTULIN KD HaCaTs left untreated or treated with proteasome inhibitor MG132 for 16 hrs. α-actin is shown as loading control. **d.** Co-immunoprecipitation (IP) using an antibody against β-catenin on lysates from WT and OTULIN KD HaCaTs left untreated or treated with proteasome inhibitor MG132 for 16 hrs. IP lysates were untreated or supplemented with recombinant human OTULIN post-IP. α-actin is shown as loading control. **e.** Co-immunoprecipitation (IP) using an antibody against β-catenin on lysates from PMK cultures isolated from OTULIN^FL/FL^ and Δ^Ker^OTULIN mice left untreated or treated with proteasome inhibitor MG132 for 16 hrs. α-actin is shown as loading control. **f.** K48 Ubq pulldown by TUBE-IP on PMKs cultures isolated from OTULIN^FL/FL^ and Δ^Ker^OTULIN mice left untreated or treated with proteasome inhibitor MG132 for 16 hrs, followed by immunoblotting for K48 ubiquitin chains and total β-catenin. Pre-immunoprecipitation (PIP) lysates were blotted for the same proteins and α-actin is shown as loading control.

### OTULIN-dependent Wnt signalling protects keratinocytes from death

We and others previously demonstrated that skin inflammation in Δ^Ker^OTULIN mice is driven by TNF-mediated apoptotic and necroptotic KC death^3,4^. In agreement, we demonstrated that OTULIN-deficient PMKs exhibit heightened sensitivity to cell death^3,4^. To investigate whether the enhanced sensitivity of OTULIN-deficient KCs to TNF-induced cell death is normalised upon β-catenin stabilisation, we stimulated PMKs isolated from Δ^Ker^OTULIN, Δ^Ker^OTULIN ΔNβ-cat and control mice with TNF and IFNγ. OTULIN-deficient KCs with stabilised β-catenin are markedly protected from TNF-induced apoptosis and necroptosis, as evidenced by decreased levels of cleaved caspase-3 and phosphorylated MLKL (pMLKL) compared to Δ^Ker^OTULIN KCs (Figure 5a). Notably, reduced levels of TNF-induced cell death were also observed in PMKs isolated from OTULIN-proficient ΔNβ-cat mice relative to cells from control OTULIN^fl/fl^ littermates, indicating that enhanced canonical Wnt signalling increases the threshold for TNF toxicity in adult KCs (Figure 5a). *In vivo*, reduced levels of cleaved caspase-8 and cleaved caspase-3 were detected in tail wholemounts from Δ^Ker^OTULIN ΔNβcat compared to Δ^Ker^OTULIN epidermis (Figure 5b). To analyse whether the increased KC cell death levels induce Wnt repression in Δ^Ker^OTULIN skin, we quantified Wnt target gene expression in Δ^Ker^OTULIN/FADD/MLKL mice, which are protected from FADD and MLKL-dependent KC death and subsequent skin inflammation^3^. Quantitative PCR analyses indicated that the levels of important Wnt mediators and target genes were similarly reduced in both Δ^Ker^OTULIN/FADD/MLKL and Δ^Ker^OTULIN skin (Figure 5c), indicating that OTULIN promotes Wnt signalling independently of KC cell death levels. Collectively, these findings indicate that the induction of Wnt signalling by OTULIN sets the threshold for KC susceptibility to TNF-induced apoptosis and necroptosis.

**Figure 5.**
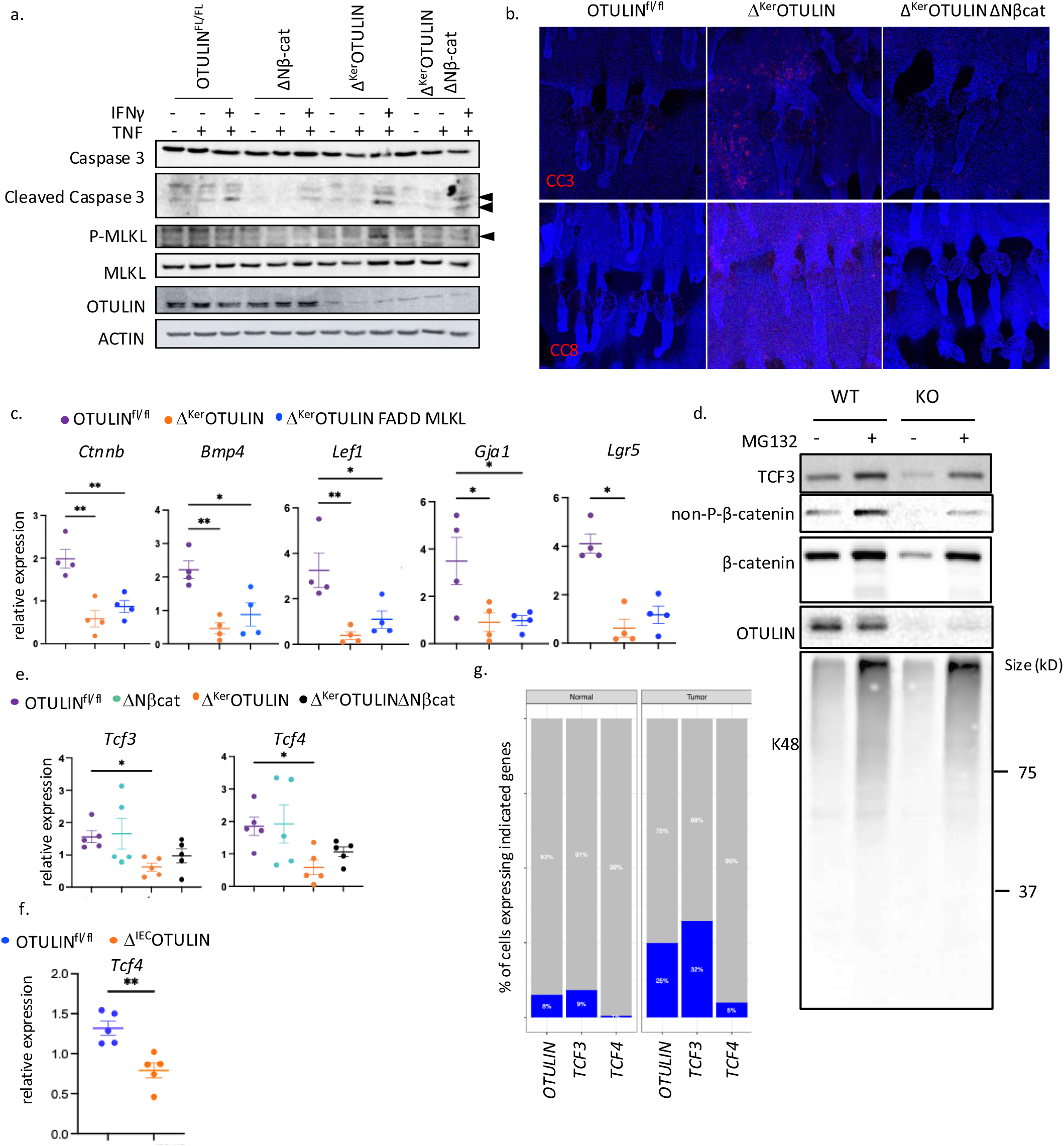
OTULIN-driven Wnt signalling protects KCs from death. **a.** Immunoblotting of lysates from PMK cultures isolated from OTULIN^FL/FL^ and Δ^Ker^OTULIN mice with antibodies against (cleaved) caspase-3, (phosphorylated) MLKL and OTULIN. Cells were untreated, treated with TNF or pretreated overnight with IFNγ prior to TNF treatment. Actin blot is shown as loading control. Arrowheads indicate cleavage fragments of caspase-3 and p-MLKL respectively. **b.** Immunofluorescent staining for cleaved caspase-3 (CC3) and cleaved caspase-8 (CC8) on epidermal tail wholemounts from OTULIN^FL/FL^, Δ^Ker^OTULIN and Δ^Ker^OTULIN ΔNβ-cat mice. **c.** Relative mRNA expression of the indicated Wnt target genes in epidermal tail lysates of OTULIN^FL/FL^, Δ^Ker^OTULIN and Δ^Ker^OTULIN FADD MLKL mice. Data represent means ± SEM (*p< 0.05; **p < 0.01; ***p < 0.001). **d.** Western blot analysis of lysates from untreated or MG132-treated PMK cultures isolated from OTULIN^FL/FL^ and Δ^Ker^OTULIN mice using antibodies against TCF3, β-catenin, non-phosphorylated β-catenin, OTULIN and K48 ubiquitin. **e.** Relative mRNA expression of *Tcf3* and *Tcf4* in epidermal tail lysates of OTULIN^FL/FL^, Δ^Ker^OTULIN and Δ^Ker^OTULIN ΔNβcat mice. Data represent means ± SEM (*p< 0.05). **f.** Relative mRNA expression of *Tcf4* in intestinal lysates of OTULIN^FL/FL^ and Δ^IEC^OTULIN mice. Data represent means ± SEM (p= 0.0035). **g**. Percentage of epithelial cells expressing OTULIN, TCF3 or TCF4 in normal human colon tissue versus colorectal cancer (tumour) samples as observed by scRNAseq^36^.

### OTULIN regulates the stability of the survival factors TCF3 and TCF4

To explore how linear ubiquitination of β-catenin influences KC survival, we focused on two Wnt target genes, *Tcf3* and *Tcf4*, which were previously shown to protect cells from death by cooperative functioning with β-catenin to drive transcription^6,23,32^*. Tcf3* and *Tcf4* were significantly suppressed in OTULIN-deficient KCs (Figure 1c, d). Previous studies demonstrated that genetic ablation of *Tcf3/4* in KCs results in a marked increase of dying cells in neonatal mouse epidermis^6^. Immunostaining confirmed expression of TCF4 in adult normal HFs^33^, whereas no staining was detectable in Δ^Ker^OTULIN skin (Supplementary Figure 4a). Immunoblotting for TCF3 revealed a reduction in protein expression in PMKs isolated from Δ^Ker^OTULIN relative to control cells (Figure 5d). Proteasome inhibition stabilised TCF3 in both wild-type and knockout cells, indicating that TCF3 stability is regulated by K48 ubiquitination in KCs. UBAN-IP on lysates obtained from wild-type and OTULIN-deficient PMKs indicated that TCF3 is not pulled down in these cells, indicating that TCF3 is not linearly ubiquitinated in cultured mouse KCs (Supplementary Figure 4b). Both Wnt-repressive and -activating properties have been attributed to TCF3/4^6^. In this context, TCF3 expression in PMKs strongly correlated with the levels of activated β-catenin, suggesting that TCF3 functions as a Wnt activator in these conditions (Figure 5d). Consistently, *Tcf3* and *Tcf4* levels were reduced in epidermal lysates obtained from Δ^Ker^OTULIN skin and normalised in epidermis from Δ^Ker^OTULIN ΔNβcat mice (Figure 5e). These findings suggest that the protection from cell death observed in KCs from Δ^Ker^OTULIN ΔNβcat mice could be attributed to enhanced TCF3/4 activity.

Finally, we questioned whether OTULIN could drive canonical Wnt signalling in tissues beyond the skin. Indeed, TCF3/4 are transcription factors that are associated with repression of differentiation, not only in skin, but also in intestinal epithelia^32,34^. Analysis of *Tcf4* levels in intestinal epithelial cells (IECs) isolated from IEC-selective OTULIN-deficient and wild-type mice confirmed a significant downregulation of *Tcf4* in OTULIN-deficient IECs (Figure 5f). Supporting a role for OTULIN in regulating stemness, our previously published scRNAseq dataset comparing control and Δ^Ker^OTULIN non-lesional (NL) and lesional (L) skin^3^ revealed higher expression of the HF stem cell markers *Cd34* and *Lgr5* in OTULIN-positive versus OTULIN-negative KCs (Supplementary Figure 4c). Given the frequent dysregulation of Wnt signalling in cancer and the dependence of various tumour types on canonical Wnt activation for growth and progression^23^, these findings suggest a broader role for OTULIN in modulating Wnt-driven processes. Colorectal cancer is a well-characterised Wnt-driven tumour type, with progression often promoted by mutations in Wnt pathway genes such as *APC*, *β-catenin* and *AXIN2*^35^. Analysis of a publicly available scRNAseq dataset of human colorectal cancer samples and healthy colon tissue (Human colon cancer atlas, GEO: GSE178341)^36^, revealed elevated levels of *OTULIN, TCF3* and *TCF4* in epithelial tumour cells compared to normal intestinal epithelial cells (Figure 5g, Supplementary Figure 4d). These findings suggest that OTULIN-mediated promotion of Wnt signalling may play a role in colorectal cancer. Together, these results highlight OTULIN as a driver of Wnt signalling across various epithelial tissues and disease contexts.

## Discussion

Protection from cell death is crucial in KCs to ensure proper epidermal barrier function. The physiological impact of KC death depends on the mode of cell death and the location of their death^37^. Aberrant linear deubiquitination due to absence of OTULIN, primes KCs for death by apoptosis and necroptosis, processes that drive dermatitis and squamous carcinogenesis^3,4,38^. In this study, we present physiological evidence positioning OTULIN as a dual-function survival factor in KCS. We show that OTULIN’s DUB activity not only stabilises LUBAC to prevent TNF-induced cell death and inflammation, but also promotes canonical Wnt signalling in KCs. Notably, preventing cell death by depletion of cell death executioners in OTULIN-deficient KCs failed to restore Wnt target gene expression, indicating that Wnt downregulation occurs independently of KC death. Stabilisation of β-catenin in KCs, through genetic crossing with mice expressing a truncated version of β-catenin, potently halts or prevents progressive dermatitis caused by OTULIN-deficiency. This truncated version of β-catenin lacks the lysine residues required for K48 ubiquitination, and K133, the site implicated in linear ubiquitination by LUBAC^27^. Induction of truncated β-catenin expression in KCs of Δ^Ker^OTULIN mice significantly reduced KC death and subsequent skin inflammation. The prosurvival effects of β-catenin stabilisation were independent of LUBAC protein levels, which remained downregulated in Δ^Ker^OTULIN ΔNβcat mice. These data indicate that canonical Wnt signalling can, at least partially, bypass cell death checkpoints controlled by LUBAC activity^39^. Strikingly, PMKs from Δ^Ker^OTULIN ΔNβcat mice exhibited reduced TNF-induced apoptotic and necroptotic cell death relative to KCs from Δ^Ker^OTULIN mice. The TNF-detoxifying effects of stabilised β-catenin were also observed in OTULIN-proficient KCs, suggesting that the extent of canonical Wnt signalling sets the threshold for TNF toxicity in KCs.

Previous studies have suggested a role for OTULIN in stabilising β-catenin in breast cancer cell lines exposed to genotoxic stress^27^. Our findings extend this concept, demonstrating that β-catenin undergoes extensive linear ubiquitination in normal adult KCs both *in vivo* and *in vitro*. In absence of OTULIN, β-catenin exhibits increased K48-linked ubiquitination, leading to its proteasomal degradation. We establish that this regulatory mechanism operates in homeostatic KCs, where β-catenin interacts with OTULIN to modulate KC susceptibility to cell death by apoptosis and necroptosis. Supporting this, a study probing for linearly ubiquitinated substrates in glioblastoma cells identified β-catenin as an M1 ubiquitination target^40^. However, it remains unclear whether OTULIN-driven Wnt signalling is a general mechanism to safeguard cell viability in conditions of stress.

The levels of activated β-catenin closely correlated with TCF3 expression in KCs and both were markedly reduced in absence of OTULIN. TCF3/4 act as transcription factors upon binding with β-catenin^32^ and are essential regulators of postnatal KC survival^6^. We show that TCF3 is not linearly ubiquitinated in KCs, but its stability appears to be regulated by forming a complex with β-catenin, which is degraded in OTULIN-deficient KCs. Notably, TCF3/4 are known to repress differentiation in postnatal skin and intestinal epithelial cells^6,33,34^. In OTULIN-deficient KCs, TCF3/4 are markedly downregulated, potentially opposing differentiation of stem cells into HF cells. This is supported by our scRNAseq data, which reveals a marked increase in the relative abundance of HF bulge cells in Δ^Ker^OTULIN skin. In intestinal epithelial cells, TCF3/4 levels are downregulated in adult cells, but are re-expressed in epithelial colorectal tumour cells. This re-expression is linked to a more undifferentiated state and coincides with OTULIN upregulation. Together, these findings indicate that OTULIN driving canonical Wnt signalling and thereby TCF3/4 expression, may constitute a survival pathway used by various cell-types in diverse disease contexts.

In conclusion, our findings establish OTULIN as a critical regulator of KC survival by performing a dual role: mediating the linear (de)ubiquitination of proteins involved in the TNF pathway, and inducing canonical Wnt signalling upstream of cell death. This two-tiered mechanism likely acts as a safeguard to suppress differentiation in conditions of heightened proliferative needs, as is the case in inflammation.

## Methods

## Acknowledgements

We thank Fiona Watt for kindly providing K14ΔNβcatenin mice. We thank the VIB Bioimaging Core and VIB Flow cytometry Core for assistance with experiments and the animal caretakers of the Center for Inflammation Research for colony maintenance of the mice. Research in the van Loo lab is supported by VIB and research grants from Ghent University (BOF23/GOA/001), FWO (3G090322 and 3G0H2522), the Charcot Foundation, the Queen Elisabeth Medical Foundation and the FOREUM Foundation. This research was funded by FWO research projects (3G032320 and G071224N), Foundation against cancer (365L04523), LEO Foundation Award Region EMEA 2022 and the Special Research Fund of Ghent University (BOF/24J/2023/138).

## Author contributions

K.L. performed most of the experiments, analysed data and drafted the manuscript. P.H. performed experiments and analysed data. A.T. and F.B. and L.V. performed experiments and analysed data. M.C. carried out bioinformatics analyses on scRNAseq data. K.S. helped in generating OTULIN-deficient HaCaTs. GvL provided critical scientific input on experiment design and the manuscript. E.H. designed and performed experiments, analysed data and drafted the manuscript.

## Supplementary figure legends

**Supplementary Figure 1.** Induction of canonical Wnt signalling in keratinocytes. **a.** Heatmap showing the expression of the indicated genes in scRNAseq data from KCs isolated from OTULIN^FL/FL^ and Δ^Ker^OTULIN skin. Each line represents a single KC. **b.** Western blot analysis of the unbound fraction and on full epidermal tail lysates from OTULIN^FL/FL^ and Δ^Ker^OTULIN mice. Concanavalin A-Sepharose beads were used to capture glycosylated E-cadherin at adherens junctions that associate with β-catenin. α-actin blot is shown as loading control.

**Supplementary Figure 2.** Constitutive activation of canonical Wnt signalling protects Δ^Ker^OTULIN mice from dermatitis. **a.** Representative pictures of mouse back and tail skin of Δ^Ker^OTULIN ΔNβ-cat mice at the age of 40 weeks. **b.** H&E-stained skin sections of 11 week-old Δ^Ker^OTULIN and Δ^Ker^OTULIN ΔNβ-cat mice. Scale bars: 200 μm. **c.** Epidermal thickness measured on H&E-stained skin sections (NL: non-lesional; L: lesional) from tamoxifen-treated Δ^Ker^OTULIN and Δ^Ker^OTULIN ΔNβ-cat 11 week-old mice (n≥ 4 mice per condition, for each biological replicate the mean of 10 measurements was taken; \*\*\*\**p* < 0.0001; two-way ANOVA with multiple comparisons). **d.** Immunoblotting for OTULIN, SHARPIN and HOIP in epidermal tail lysates from OTULIN^FL/FL^, ΔNβ-cat, Δ^Ker^OTULIN and Δ^Ker^OTULIN ΔNβ-cat mice. α-actin is shown as loading control.

**Supplementary Figure 3.** Generation of OTULIN-deficient human KCs and immunoprecipitation studies in mouse and human KCs. **a.** Quantitative spectrum of indels created by two consecutive treatments with OTULIN-targetting nanoblades as assessed by TIDE (Tracking of Indels by DEcomposition) analysis. **b.** Immunoblot for OTULIN on lysates from wild-type (WT) and OTULIN knock-down (OTULIN KD) HaCaTs. α-actin is shown as loading control. **c.** WT and OTULIN KD HaCaTs (n=3 technical replicates per condition) were treated with 30 ng/ml hTNF for 24 hrs. Viability was assessed by Sytox Green uptake. Data represent means ± SEM. (*p<0.05; REML analysis). **d.** Full blots of the immunoprecipitation experiments shown in Figure 4c, including visualization of control (beads only, no IP antibody). Asterisk depicts an aspecific band; arrowhead indicates OTULIN-specific band.

**Supplementary Figure 4.** OTULIN regulates TCF3/4 levels in health and disease. **a.** Immunofluorescent staining for TCF4 in epidermal tail wholemounts of OTULIN^FL/FL^ and Δ^Ker^OTULIN mice. **b.** Linear Ubq pulldown by UBAN-immunoprecipitation (IP) on PMK cultures isolated from OTULIN^FL/FL^ and Δ^Ker^OTULIN mice and left untreated or treated with MG132 for 8 hrs, followed by immunoblotting for M1 ubiquitin chains, β-catenin and TCF3. **c.** Feature plots showing expression levels of *OTULIN* in epithelial cells from normal colon (left) and in tumour epithelial cells from colorectal cancer (right) as assessed by scRNAseq^36^. **d.** Violin plots showing the normalised expression levels of hair follicle stem cell markers *Cd34* and *Lgr5* in OTULIN positive KCs versus OTULIN negative KCs.

