## Supplementary FIgure 1 for "Canonical Wnt induction by OTULIN prevents keratinocyte death and skin inflammation"

a. b.

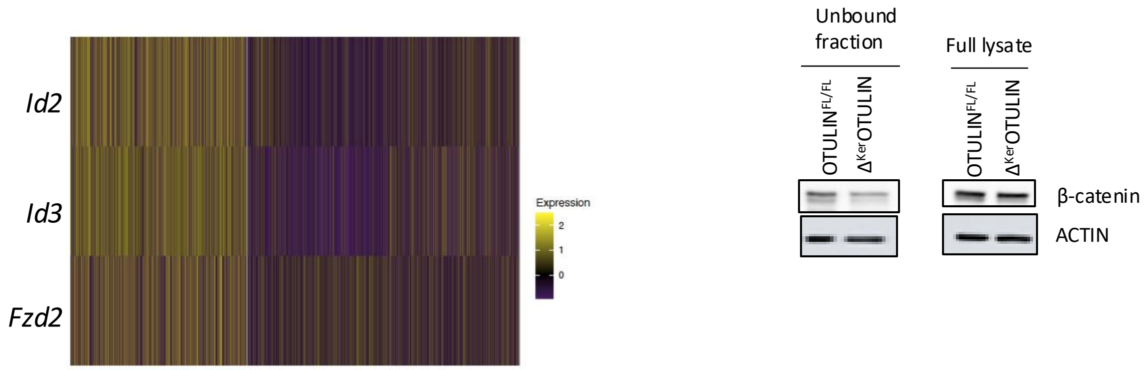


**Supplementary Figure 1. Induction of canonical Wnt signalling in keratinocytes. a.** Heatmap showing the expression of the indicated genes in scRNAseq data from KCs isolated from OTULIN$^{FL/FL}$ and $\Delta^{Ker}$OTULIN skin. Each line represents a single KC. **b.** Western blot analysis of the unbound fraction and on full epidermal tail lysates from OTULIN$^{FL/FL}$ and $\Delta^{Ker}$OTULIN mice. Concanavalin A-Sepharose beads were used to capture glycosylated E-cadherin at adherens junctions that associate with β-catenin. α-actin blot is shown as loading control.
