## Supplementary FIgure 2 for "Canonical Wnt induction by OTULIN prevents keratinocyte death and skin inflammation"

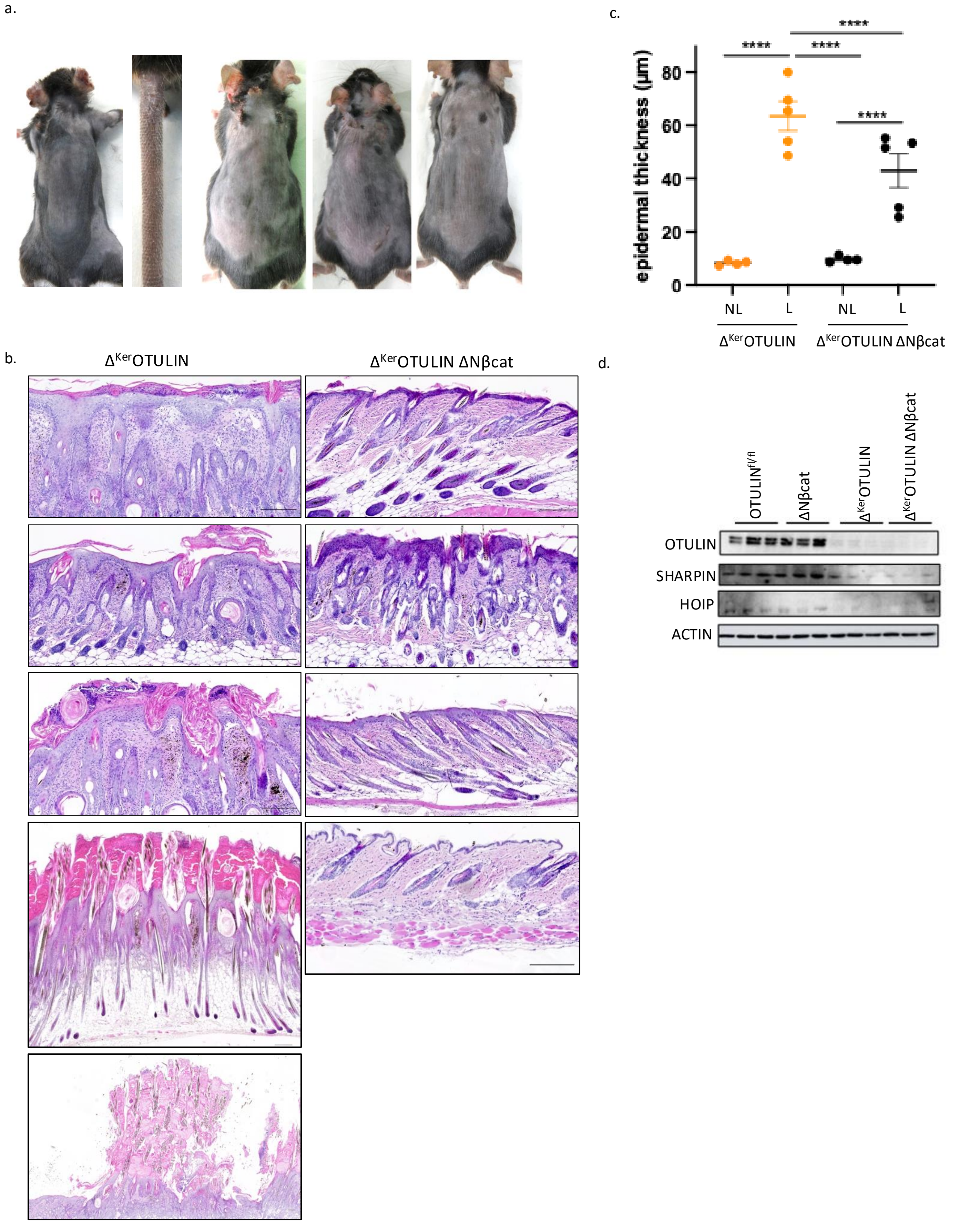


**Supplementary Figure 2. Constitutive activation of canonical Wnt signalling protects $\Delta^{Ker}$OTULIN mice from dermatitis. a.** Representative pictures of mouse back and tail skin of $\Delta^{Ker}$OTULIN ΔNβ-cat mice at the age of 40 weeks. **b.** H&E-stained skin sections of 11 week old $\Delta^{Ker}$OTULIN and $\Delta^{Ker}$OTULIN ΔNβ-cat mice. Scale bars: 200 µm. **c.** Epidermal thickness measured on H&E-stained skin sections (NL: non-lesional; L: lesional) from tamoxifen-treated $\Delta^{Ker}$OTULIN and $\Delta^{Ker}$OTULIN ΔNβ-cat 11 week-old mice (n≥ 4 mice per condition, for each biological replicate the mean of 10 measurements was taken; $^{****}p < 0.0001$; two-way ANOVA with multiple comparisons). **d.** Immunoblotting for OTULIN, SHARPIN and HOIP in epidermal tail lysates from $OTULIN^{FL/FL}$, ΔNβ-cat, $\Delta^{Ker}$OTULIN and $\Delta^{Ker}$OTULIN ΔNβ-cat mice α-actin is shown as loading control.
