## Supplementary FIgure 3 for "Canonical Wnt induction by OTULIN prevents keratinocyte death and skin inflammation"

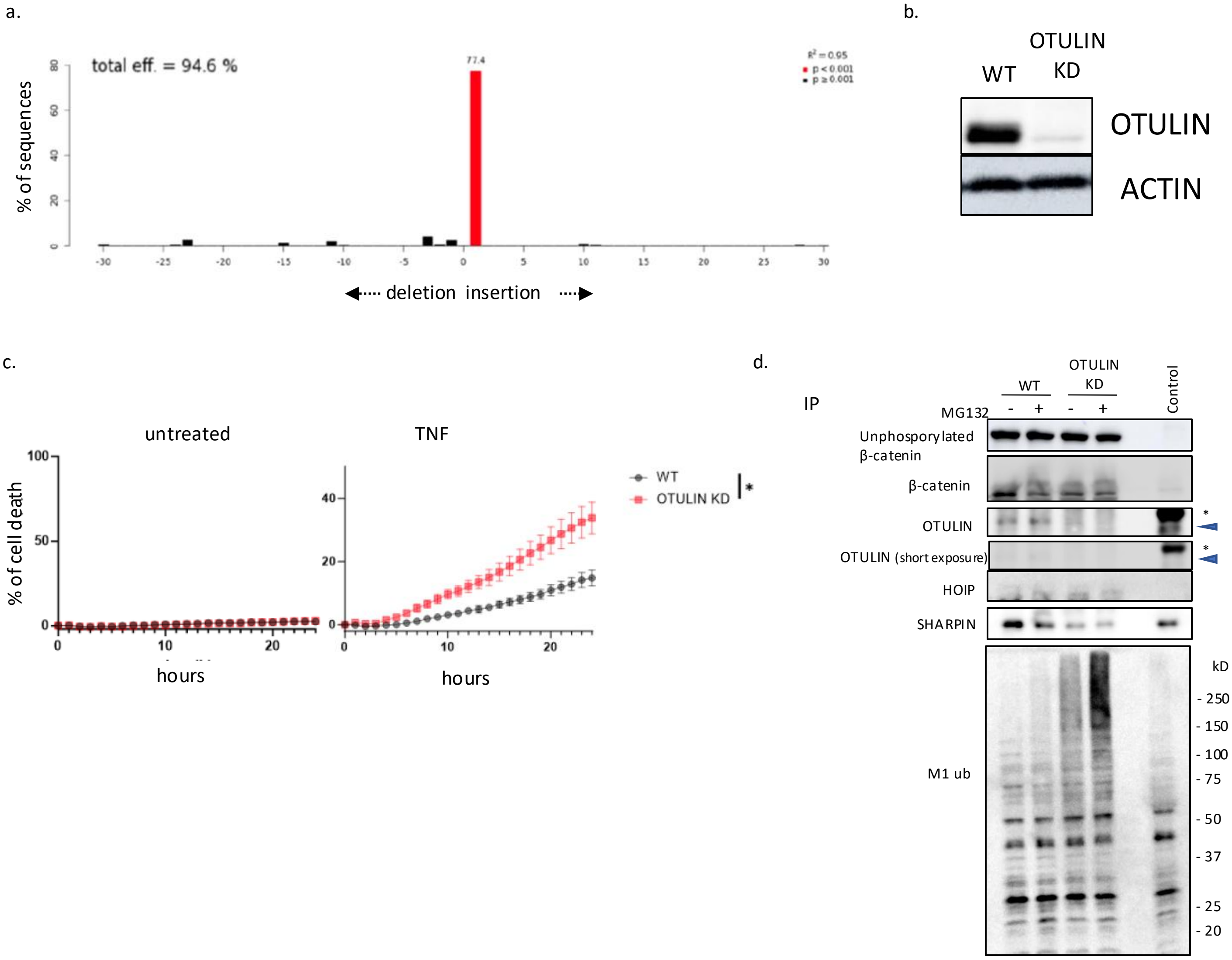


**Supplementary Figure 3. Generation of OTULIN-deficient human KCs and immunoprecipitation studies in mouse and human KCs. a.** Quantitative spectrum of indels created by two consecutive treatments with OTULIN-targetting nanoblades as assessed by TIDE (Tracking of Indels by DEcomposition) analysis. **b.** Immunoblot for OTULIN on lysates from wild-type (WT) and OTULIN knock-down (OTULIN KD) HaCaTs. $\alpha$-actin is shown as loading control. **c.** WT and OTULIN KD HaCaTs (n=3 technical replicates per condition) were treated with 30 ng/ml hTNF for 24 hrs. Viability was assessed by Sytox Green uptake. Data represent means ± SEM. (*p<0.05; REML analysis). **d.** Full blots of the immuno-precipitation experiments shown in Figure 4c, including visualization of control (beads only, no IP antibody). Asterisk depicts an aspecific band; arrowhead indicates OTULIN-specific band.
