## Supplementary FIgure 4 for "Canonical Wnt induction by OTULIN prevents keratinocyte death and skin inflammation"

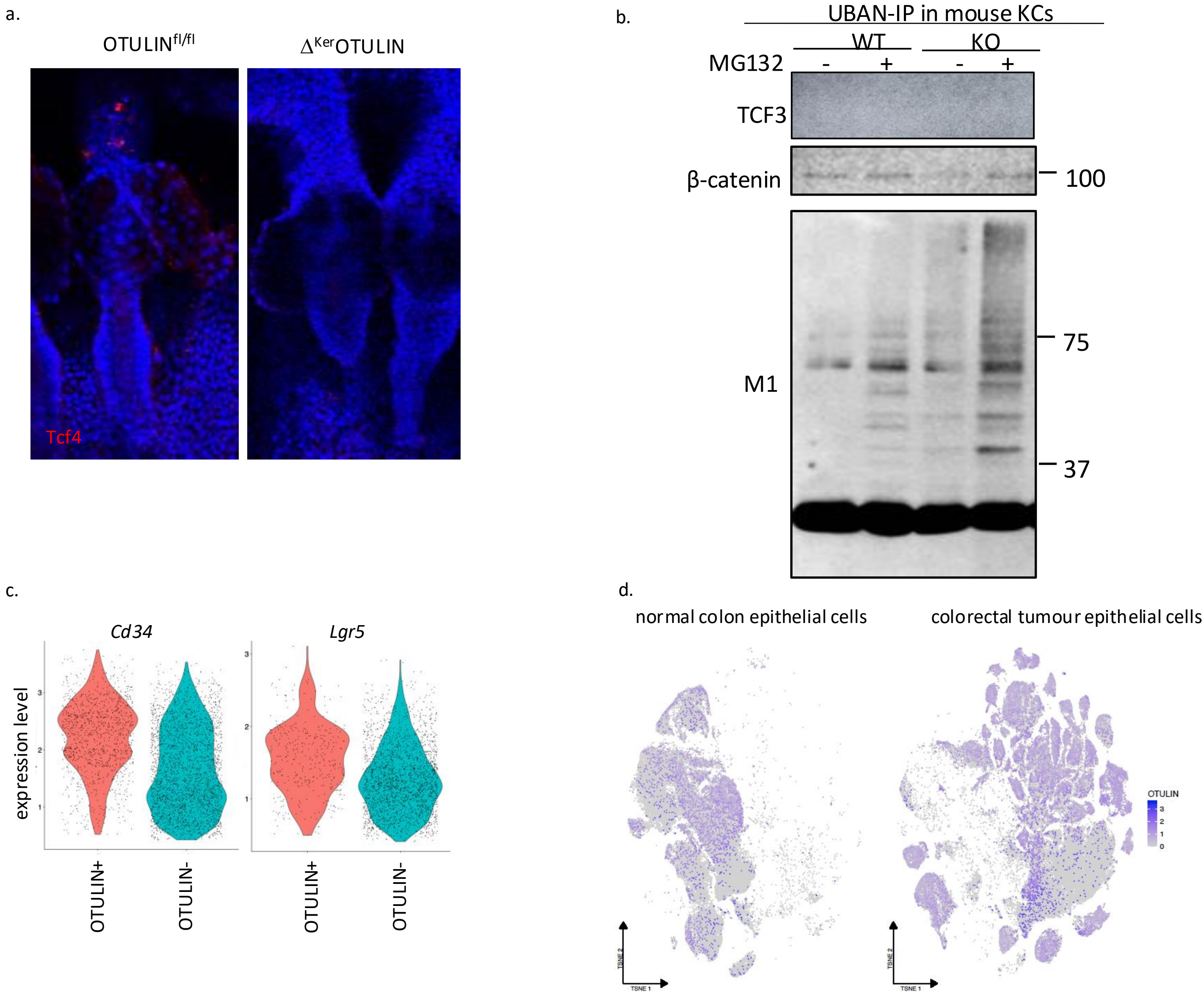


**Supplementary Figure 4. OTULIN regulates TCF3/4 levels in health and disease. a.** Immunofluorescent staining for TCF4 in epidermal tail wholemounts of $OTULIN^{FL/FL}$ and $\Delta^{Ker}OTULIN$ mice. **b.** Linear Ubq pulldown by UBAN-immuno-precipitation (IP) on PMK cultures isolated from $OTULIN^{FL/FL}$ and $\Delta^{Ker}OTULIN$ mice and left untreated or treated with MG132 for 8 hrs, followed by immunoblotting for M1 ubiquitin chains, β-catenin and TCF3. **c.** Feature plots showing expression levels of *OTULIN* in epithelial cells from normal colon (left) and in tumour epithelial cells from colorectal cancer (right) as assessed by scRNAseq[36]. **d.** Violin plots showing the normalised expression levels of hair follicle stem cell markers *Cd34* and *Lgr5* in OTULIN positive KCs versus OTULIN negative KCs.
